# A genetic bottleneck of mitochondrial DNA during human lymphocyte development

**DOI:** 10.1101/2021.09.28.462268

**Authors:** Zhongjie Tang, Zhaolian Lu, Baizhen Chen, Weixing Zhang, Howard Y. Chang, Zheng Hu, Jin Xu

**Affiliations:** State Key Laboratory of Biocontrol, School of Life Sciences, Sun Yat-Sen University, Guangzhou, China; CAS Key Laboratory of Quantitative Engineering Biology, Shenzhen Institute of Synthetic Biology, Shenzhen Institutes of Advanced Technology, Chinese Academy of Sciences, Shenzhen 518055, China; Center for Personal Dynamic Regulomes, Stanford University, United States; Departments of Dermatology and Genetics, Stanford University School of Medicine, United States; Howard Hughes Medical Institute, Stanford University, United States

## Abstract

Mitochondria are essential organelles in eukaryotic cells that provide critical support for energetic and metabolic homeostasis. Mutations that accumulate in mitochondrial DNA (mtDNA) in somatic cells have been implicated in cancer, degenerative diseases, and the aging process. However, the mechanisms used by somatic cells to maintain proper functions despite their mtDNA mutation load are poorly understood. Here, we analyzed somatic mtDNA mutations in more than 30,000 human single peripheral and bone marrow mononuclear cells and observed a significant overrepresentation of homoplastic mtDNA mutations in B, T and NK lymphocytes despite their lower mutational burden than other hematopoietic cells. The characteristic mutational landscape of mtDNA in lymphocytes were validated with data from multiple platforms and individuals. Single-cell RNA-seq and computational modeling demonstrated a stringent mitochondrial bottleneck during lymphocyte development likely caused by lagging mtDNA replication relative to cell proliferation. These results illuminate a potential mechanism used by highly metabolically active immune cells for quality control of their mitochondrial genomes.

## INTRODUCTION

Mitochondrial DNA (mtDNA) encodes genes involved in oxidative phosphorylation that are essential for eukaryotic cells^1^. There are typically hundreds to thousands of copies of mtDNA molecules in each cell and the germline mtDNA is predominantly maternally inherited and does not undergo recombination^2^. mtDNA accumulates mutations at a rate that is five to ten times higher per site than the nuclear genome because the lack of DNA repair systems^3,4^ and frequent contact with mutagenic reactive oxygen species (ROS)^5^. More than 500 pathogenic mtDNA mutations have been identified as causative genetic defects of various human diseases^6^. According to the theory known as “Muller’s ratchet,” continuous accumulation of deleterious mutations in the absence of purifying selection will lead to a decline in population fitness and will ultimately result in mutational meltdown^7^. To avoid this outcome, the animal germline has evolved a mitochondrial genetic bottleneck, wherein only a small subset of mtDNA is transmitted to the next generation, thus resulting in significant removal of deleterious mutations^8-10^. Population studies have also revealed an increase in mtDNA heteroplasmy in blood cells as part of the normal aging process^11^ and the accumulation of pathogenic mtDNA mutations has been reported in cancers and neurodegenerative disorders^12,13^. However, the transmission and clonal dynamics of somatic mtDNA mutations along tissue development are largely unknown, due to the technical difficulties of detecting heteroplasmic mutations in single cells.

We and others have recently developed a single-cell lineage tracing method leveraging the somatic mtDNA mutations detected in single-cell assay for transposase-accessible chromatin with high-throughput sequencing (scATAC-seq) and/or RNA-seq (scRNA-seq) data^14,15^. Using this method, a recent study had shown that a pathogenic mutation 3243A/G, the cause of mitochondrial myopathy, encephalopathy, lactic acidosis, and stroke-like episode (MELAS)^16^, was remarkably purified in T cells as compared to other blood cells from peripheral blood mononuclear cells (PBMCs), with unknown mechanisms. These results inspired our investigation of the mtDNA mutational landscape in a large population of single cells in order to understand the clonal dynamics of mtDNA in the development of somatic cell lineages.

## RESULTS

### Somatic mutational landscape of mtDNA at single-cell resolution

In this study, we focused on human hematopoietic system where the cellular differentiation lineages have been well documented. We first identified somatic mtDNA mutations in a previously reported mitochondrial scATAC-seq (or mtscATAC-seq) dataset including more than 20,000 blood cells from a healthy 47-year-old individual^17^ (**Fig. 1a-b, Extended Data Fig. 1a, Methods**). We summarized the numbers of mutations and the variant allele frequency (VAF, also referred to as mtDNA heteroplasmic ratio) in each cell in order to compare the VAF distribution in a population of different cell types. Interestingly, we found cells of the mature lymphocyte lineages--specifically B, T, NK cells--carried a significantly lower mtDNA mutational burden as compared to those identified in hematopoietic progenitor cells, including hematopoietic stem cells (HSCs), multipotent progenitors (MPPs), lymphoid-primed multipotent progenitors (LMPPs), common lymphoid progenitors (CLPs), common myeloid progenitors (CMPs), and granulocyte-macrophage progenitors (GMPs) (**Fig. 1c and Extended Data Fig. 1b**, Wilcoxon test, *p* <2.2e^-16^). The mtDNA mutational burden was also lower in lymphocytes as compared to the myeloid and erythroid lineages (**Fig. 1c**, Wilcoxon test, *p* <2.2e^-16^). As anticipated, most somatic mtDNA mutations were detected at low VAF in individual cells in all cell types (**Fig. 1d**). However, the distribution of homoplastic mutations (i.e., those at VAF∼1) varies substantially among the different cell types. For instance, progenitor cells, including HSCs, MPPs, LMPPs, CLPs, CMPs, and GMPs, exhibit the typical monotonic decline in the number of mutations with increasing VAF (**Fig. 1d**). While this pattern was also true in both the myeloid and erythroid lineages (e.g., monocytes and erythrocytes), we observed an unanticipated increase in the number of homoplastic mutations in B, T and NK cells (**Fig. 1d**).

**Fig. 1.**
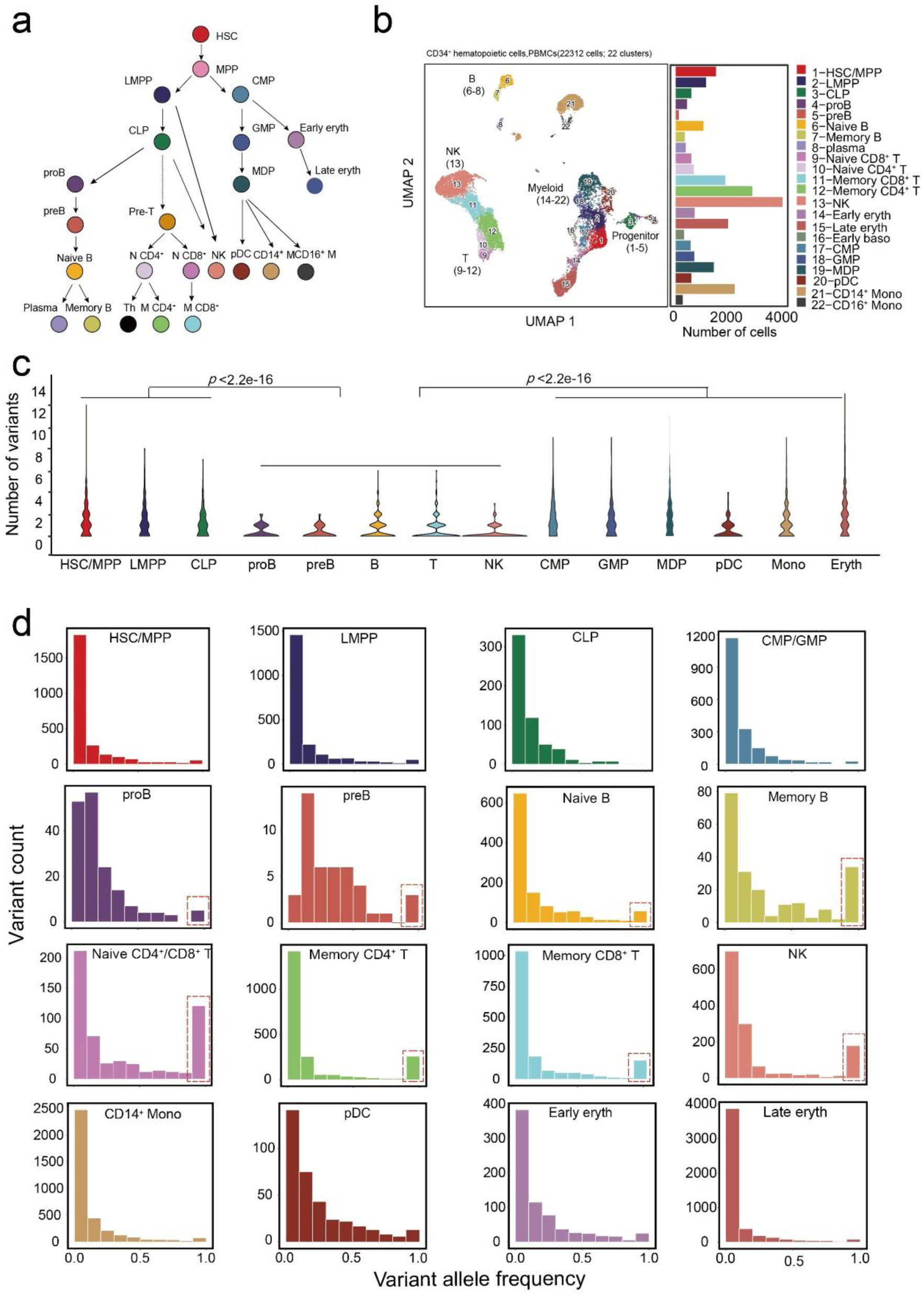
Somatic mutations in the mtDNA of PBMCs. (**a**) Schematic of human hematopoietic differentiation and lineage commitment. HSC, hematopoietic stem cell; MPP, multipotent progenitor; LMPP, lymphoid-primed multipotent progenitor; CLP, common lymphoid progenitor; CMP, common myeloid progenitor; GMP, granulocyte-monocyte progenitor; MDP, monocyte-dendritic cell progenitor; N CD4, naïve CD4^+^ T cell; N CD8, naïve CD8^+^ T cell; M CD4, memory CD4^+^ T cell; M CD8, memory CD8^+^ T cell; Th, T helper cell; NK, natural killer cell; pDC, plasmacytoid dendritic cell; Eryth, erythrocyte. (**b**) UMAP projection of 22,312 CD34+ hematopoietic cells and PBMCs with mtscATAC-seq data. Dots represent individual cells that have been colored according to cluster identity. The bar plot indicates the number of cells in each cluster (labeled at right). (**c**) Violin plot showing the number of somatic mtDNA variants per cell for various cell types; *P*-values, two-sided Wilcoxon rank-sum test. (**d**) The VAF distribution of somatic mtDNA mutations across different cell types. Homoplastic mutations (VAF ∼1) identified in the lymphoid lineage are highlighted with a red box.

In addition to the mtscATAC-seq dataset from PBMCs, we analyzed another mtscATACseq dataset of 10,327 bone marrow mononuclear cells (BMMCs) from an independent healthy donor^18^ (**Fig. 2a**). As the observations in PBMCs, lymphocytes in BMMCs also carried a lower mtDNA mutational burden with a characteristic overrepresentation of homoplastic mutations (**Fig. 2b and Extended Data Fig. 1c**). In fact, these lymphocyte-specific characteristics were also verified by additional scATAC-seq or scRNA-seq data from 7 independent individuals (**Extended Data Fig. 2**), indicating a general and unique process of clonal dynamics of mtDNA in lymphocyte development.

**Fig. 2.**
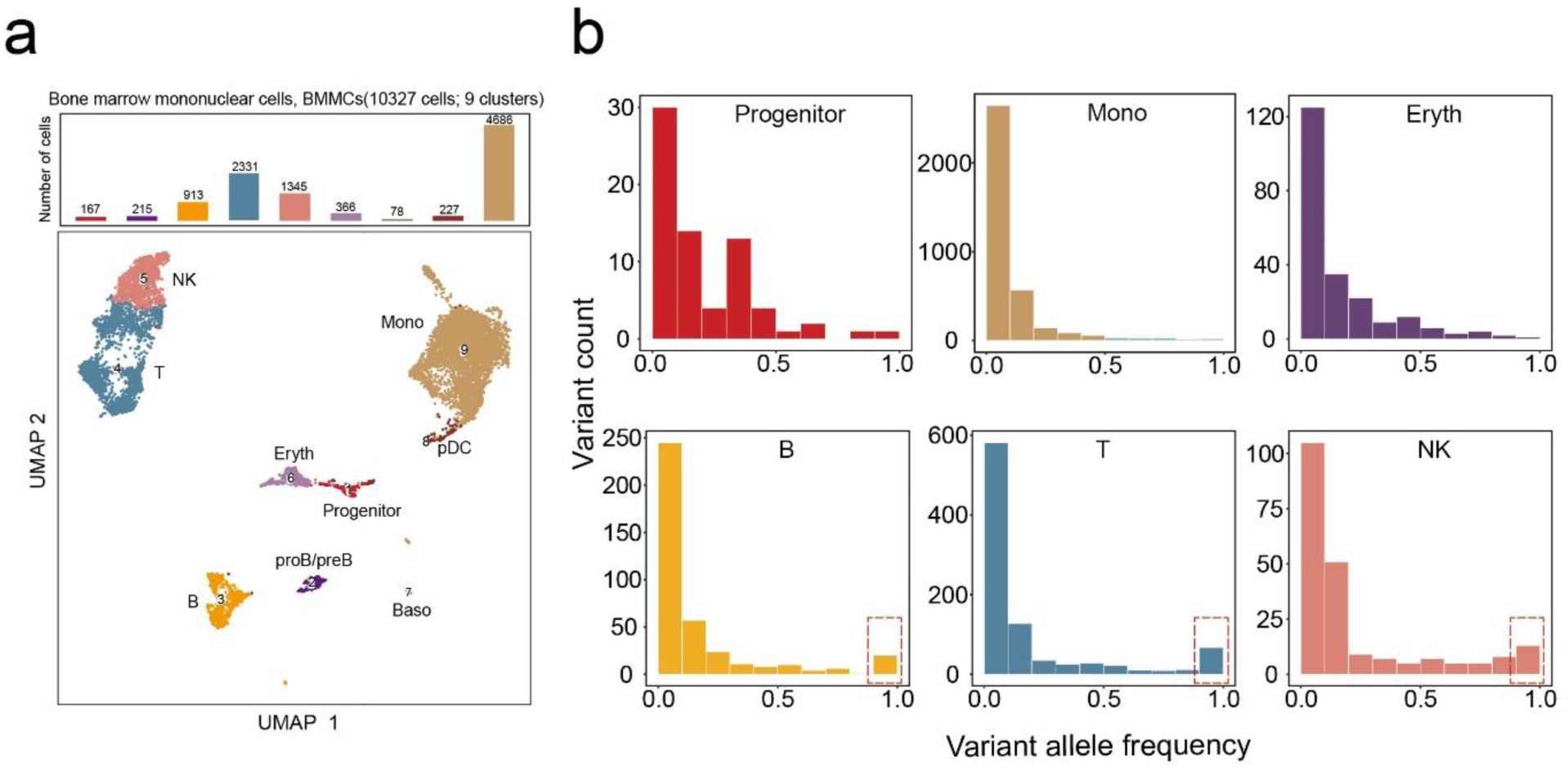
Somatic mutations in the mtDNA of BMMCs. **(a)** UMAP projection of 10,327 mononuclear cells from bone marrow with mtscATAC-seq data. Dots represent individual cells that have been colored according to cluster identify and cell types. **(b)** The VAF distribution of somatic mtDNA mutations across different cell types in BMMCs. Homoplastic mutations (VAF ∼1) identified in the lymphoid lineage are highlighted with a red box.

### Asynchronous replication of mitochondrial and nuclear genome during B cell development

To examine whether the distinct VAF distribution between lymphoid cells and myeloid/erythroid cells is due to the variation of mtDNA copy number per cell, we estimated the relative number of mtDNA copies in each cell type according to the fraction of sequencing reads mapped to the mitochondrial genome relative to the total number of reads in each cell (**Fig. 3a and Extended Data Fig. 3a**). Although mature lymphocytes and progenitor cells had similar mtDNA copy numbers, pro-B and pre-B cells—the earliest lineage-committed cells in B cell development—exhibited a significantly lower number of mtDNA copies (Wilcoxon test, pro-B/pre-B versus HSC/MPP, *p* <2.2e^-16^; pro-B/pre-B versus B, *p* <2.2e^-16^). Of note, the CLPs also showed significantly fewer mtDNA copies than earlier progenitors (Wilcoxon test, CLP versus HSC/MPP, *p* <2.2e^-16^; CLP versus LMPP, *p*<2.2e^-16^), thus indicating a remarkable mtDNA copy number reduction in early lymphocyte development. Therefore, we hypothesized that the characteristic mutational spectra in lymphocyte mtDNA (**Fig. 1c-d and Fig. 2b**) might result from a mitochondrial genetic bottleneck.

**Fig. 3.**
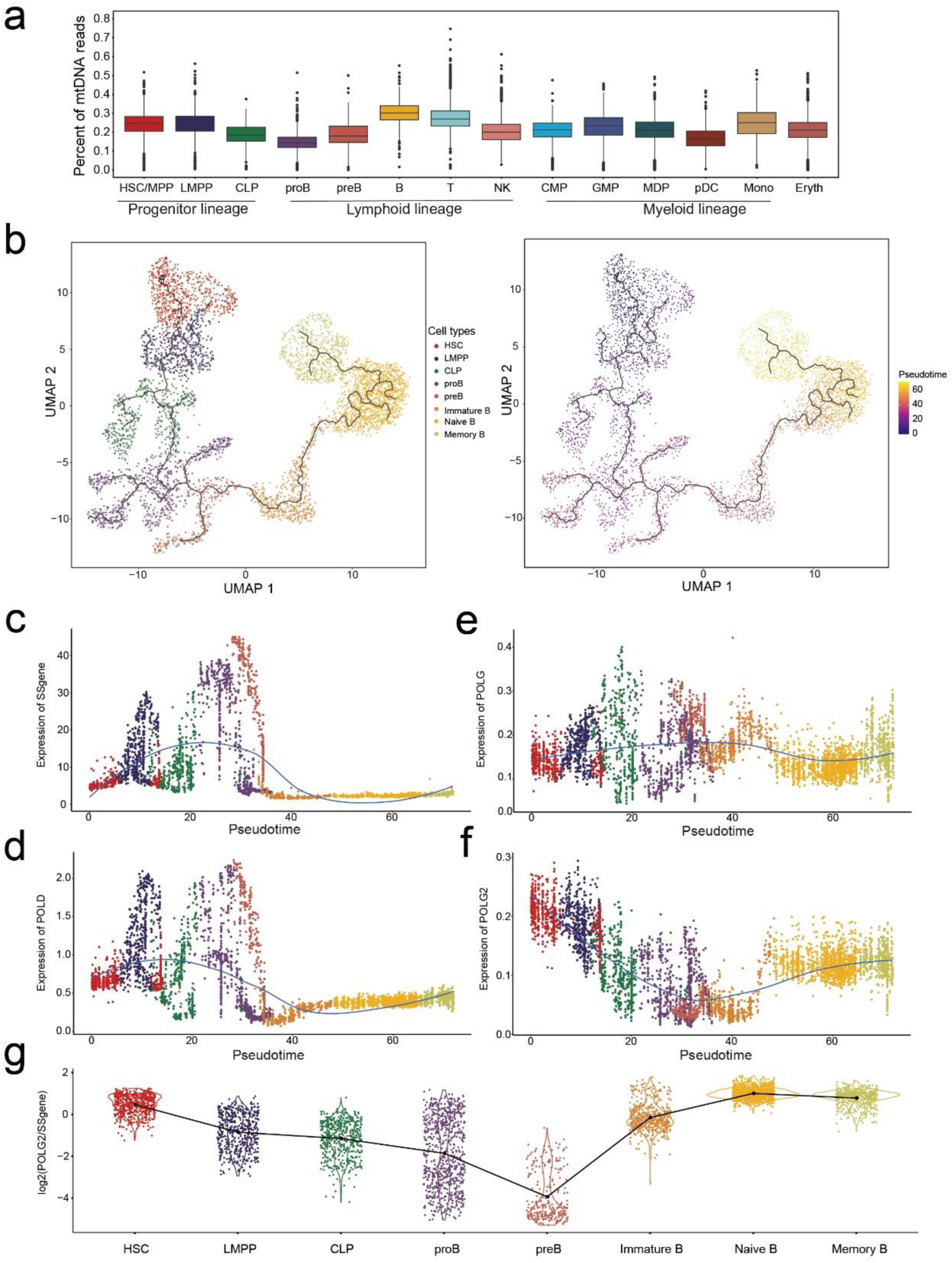
Replication of mtDNA during B cell development. **(a)** The relative number of mtDNA copies was determined by the proportion (sequencing reads mapped to mitochondrial genome divided by the total number of reads) in each cell, as identified from the scATAC-seq dataset. **(b)** Pseudo-time trajectory of B cell differentiation from HSCs by using single-cell RNA-seq data generated from PBMCs (n=4692 cells). Individual colors denote different cell types (top) and developmental stages (bottom) defined by pseudo-time. The solid line represents the fitted trajectory across pseudo-time. (**c-f**) Kinetic plots showing the expression of (**c**) 39 G1/S phase-specific genes (SSgene), (**d**) nuclear DNA polymerase δ (*POLD1–3*), (**e**) mtDNA polymerase γ (POLG) and (**f**) the binding subunit of mitochondrial DNA polymerase γ (*POLG2*) along the B-cell developmental trajectory. (**g**) Violin plot showing the ratio of *POLG2* expression to the mean expression of all G1/S phase-specific genes in each cell associated with B-cell development. The broken line represents the change trend of the mean ratio across different cell types.

To address this possibility, we examined the mtDNA replication machinery to gain insight into the regulation of mtDNA copy number along the lymphocyte differentiation trajectory. Since T cells matures in thymus and their progenitor, pre-T cells, are not available in by PBMCs, we focused on the B cell lineage. DNA polymerase γ is the only known mitochondrial DNA polymerase in animals^19^. DNA polymerase γ has both a catalytic (*POLG*) and a binding subunit (*POLG2*) and can catalyze the polymerization of deoxyribonucleotides. High levels of DNA polymerase γ activity have been detected in cell cycle phases S and G2 to maintain stable numbers of mtDNA during cell division^19-21^. To determine whether the expression of DNA polymerase γ increases with cell proliferation during B cell development, we projected the developmental trajectory of cell subpopulations from HSCs to mature B cells via a pseudo-time analysis with scRNA-seq data (**Fig. 3b and Extended Data Fig. 3b**). We observed up-regulation of G1/S phase-specific genes (such as DNA polymerase δ, *POLD1–3*) in both pro-B and pre-B cell populations, thus suggesting high activation of cell proliferation in these cell types (**Fig. 3c-d and Extended Data Fig. 3c-e**). In contrast, the expression of DNA polymerase γ was not coupled with cell proliferation (**Fig. 3e-f and Extended Data Fig. 3c-e**). Unexpectedly, the expression of the DNA polymerase γ binding subunit (*POLG2*) was significantly diminished in the highly proliferative pro-B and pre-B cell subpopulations (**Fig. 3g**). Together, these results imply a genetic bottleneck during B cell development which might have resulted from limited replication of mtDNA, thus diluting the mtDNA copy number throughout cell division.

### Quantification of mtDNA genetic bottleneck by computational modeling

To test our hypothesis and quantify the extent of the mitochondrial genetic bottleneck, we developed a computational model of an mtDNA dilution process based on population genetics theory (**Fig. 4a**). In this model, we assumed that only a proportion of mtDNA molecules (denoted by α) replicates during each cell cycle. This process continues for *T*_*d*_ cell cycles until the mtDNA copy number recovers to the initial levels (∼500 copies per cell estimated by Ryan et.al^22^). Using the approximate Bayesian computation (ABC) method, we estimated the model parameters for B, T and NK cell populations by using a constant mtDNA mutation rate of 10^−7^per site per cell division^23^ (**Fig. 4b and Extended Data Fig. 4a**). The model estimations showed the minimal mtDNA copy number were 21 (95% confidence interval [CI] =13–56), 13 (95% CI=12– 19) and 14 (95% CI=12–21) in each B, T and NK cell, respectively. These values were 20–40-fold lower than the normal mtDNA levels. The VAF distribution simulated with these parameter estimations recapitulated the observed data, showing a characteristic overrepresentation of homoplastic mutations (i.e., VAF∼1) and a reduced overall mutational burden (**Fig. 4c and Extended Data Fig. 4b-c**). Notably, this pattern cannot be achieved by random genetic drift alone with a constant number of mtDNA copies.

**Fig. 4.**
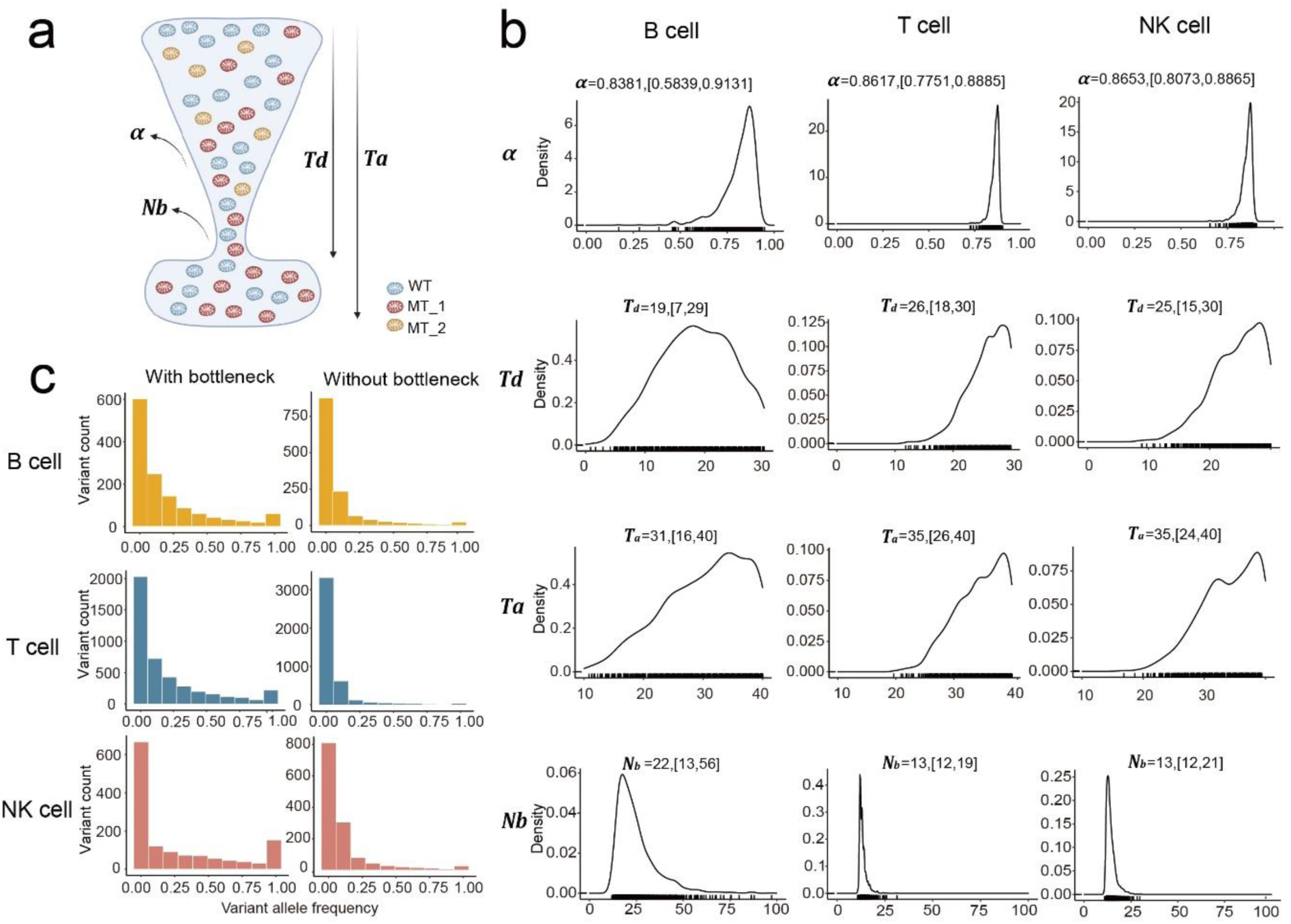
The dilution model of the mitochondrial genetic bottleneck. **(a)** Schematic illustration of the dilution model of the mitochondrial genetic bottleneck. In this model, only a fraction of mtDNA molecules (denoted by α) replicate at each cell division. After *T*_*d*_ cell divisions from the LMPP stage, the number of mtDNA copies in each lymphocyte subtype (B, T, and NK cells) undergoes rapid recovery to the baseline level (∼500 per cell). *N*_*b*_ denotes the minimal number of mtDNA copies that can be computed as Eq (1). The total number of cell divisions required for the transition from LMPP to mature lymphocyte is denoted as *T*_*a*_. **(b)** The distribution of model parameters inferred by the Approximate Bayesian Computation (ABC) algorithm. The mean and 95% confidence interval of each parameter estimation is as shown. **(c)** Simulations based on the dilution model of mitochondrial genetic bottleneck with the ABC-estimated parameter values recapitulated the lymphocyte-specific overrepresentation of homoplastic mutations and the lower mutation burden (**Extended Data Fig. 3b**). The left and right panels represent the simulations with and without mitochondrial genetic bottleneck, respectively. The average of 100 simulations carried out for each model is as shown. The results of each iteration are shown in **Extended Data Fig. 3c**.

### The consequence of mtDNA genetic bottleneck

Collectively, our integrative genomic data analysis and computational modeling demonstrated the existence of a stringent mtDNA genetic bottleneck that resulted from replicative dilution during lymphocyte development. This mechanism strengthens the genetic drift toward a lower mtDNA mutational burden and lower genetic diversity within each cell. We wondered whether the genetic bottleneck during lymphocyte development might have the same purifying selection effects as those in the germline. We thus examined the VAF distribution in various genomic regions (loop, tRNA, rRNA and coding) or mutation types (synonymous and nonsynonymous), as well as the dN/dS ratio (the ratio of the normalized number of nonsynonymous substitutions - *dN* to the normalized number of synonymous substitutions - *dS*) (**Extended Data Fig. 5a** and **Extended Data Fig. 6a**). We observed no significant differences in the VAF distribution for mutations in different genomic regions or substitution types among the various cell types. Moreover, the calculated *dN*/*dS* ratios revealed a pattern of generally neutral evolution (i.e., *dN*/*dS*∼1) in all categories in most of the cases examined (**Extended Data Fig. 5b** and **Extended Data Fig. 6b**).

Thus, our results showed that the entire mtDNA genome was evolving under a neutrality-like process. However, this is likely due to linkage of whole mitochondrial genome with strong Hill–Robertson interference, leading to a pattern of quasi-neutrality as in cancer evolution^24^. Therefore, we checked individual mutation sites to look for the signals of purifying selection and indeed observed several mutations that were specifically eliminated in lymphocytes compared to myeloid lineage (**Fig. 5a-b**). For example, the mutations, 2636G/A and 3209A/G, underwent the most profound decrease in prevalence (**Fig. 5c**) in lymphocytes. Intriguingly, these two sites are all located at MT-RNR2, which encode 16S rRNA and Humanin, a peptide playing protection roles in multiple mitochondrial diseases (**Fig. 5d**)^25^. Furthermore, we queried MITOMAP, a human mitochondrial genome database, and found that mtDNA variants reported on MT-RNR2 were highly associated with sepsis (*p*<2.2e^-16^, **Fig. 5e**)^26,27^, suggesting MT-RNR2 may play important roles in immune functions to protect from infections. These data indicate purifying selection in lymphocytes indeed occurs for specific mtDNA mutation sites.

**Fig. 5.**
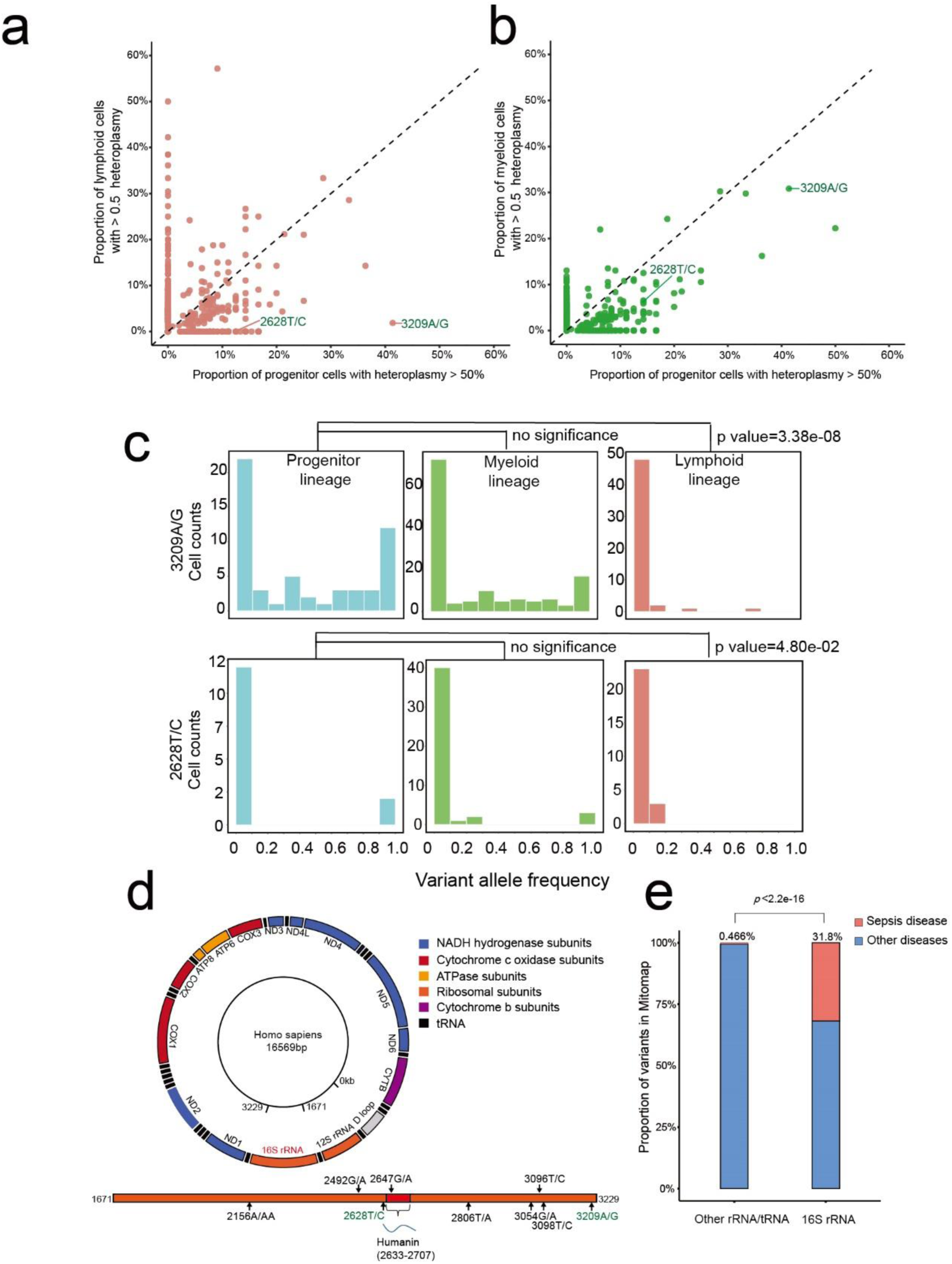
Elimination of specific mtDNA variants in lymphocyte. (**a**) Scatter plot documenting the percentage of cells with dominant mtDNA mutations (VAF>50% in a single cell). Shown are the results from progenitor cells (HSC, MPP, and LMPP) compared to cells from lymphoid (B, T, or NK cells) or myeloid lineages(**b**). **(c)** The distribution of VAF for two individual sites (3209A/G, 2636G/A) in progenitor, myeloid and lymphoid cells, respectively. The *p* values shown were determined by the Chi-square test. **(d)** The location of 16S RNA (MT-RNR2) on the mitochondrial genome and the location of sepsis association variants on MT-RNR2, reported in MITOMAP (in black), or specific eliminated in lymphocytes (in green). **(e)** The proportion of mtDNA variants associated with sepsis disease in 16S RNA versus other rRNA/tRNA genes on mitochondrial genome.

## DISCUSSION

Collectively, we observed an unanticipated lower mutational burden and accumulation of homoplastic mtDNA mutations in lymphocytes that depicted a stringent genetic bottleneck and purifying selection of mtDNA. Gene expression data and computational modeling suggest a dilution process, based on the rate of mtDNA replication relative to the nuclear genome. Although the single-cell data derived from PBMCs cannot capture the full developmental trajectory of T cells because pre-T cells develop in the thymus. Our single cell data and computational inference indicates the genetic bottleneck in T and NK cells might be as stringent as that in B cells (**Fig. 1d, Fig. 2a, Fig. 4b**). Further systematic study of T cell precursors in the thymus may provide further insight on how genetic bottleneck occurs during T cell development. Also, based on our observations and simulations, we hypothesize that the regulation of lymphocyte specific genetic bottleneck may start from CLP stage, instead of subsequent lineage commitment for B, T and NK cells. The effect of this regulation was likely enhanced via the active proliferation of progenitor cells. We knew that during lymphocyte development, multipotent T and B progenitor cells undergo a series of maturation steps that include positive selection for functional T-cell receptors (TCRs) or immunoglobulins and negative selection to eliminate cells with a high affinity for self-associated peptides or antigens^28^. Only a small proportion of T lymphoid cells will survive from the negative and positive selections. Moreover, mitochondrial function is important for T cell development and their functional activation^29,30^. The metabolic responses characteristic of lymphocytes development and activation are both well-regulated at transcriptional and post-transcriptional levels^31^. For example, several groups have shown that T or B cell activation leads to mitochondrial remodeling and dramatic shifts in cell metabolism, as part of their role in eliminating pathogens^32-36^. Meanwhile the selection against pathogenic mutations 3243 was stronger in T cells than B and NK as shown by Walker et.al^16^. All these evidences suggested that the mtDNA genetic bottleneck may be one of several potential mechanisms in the regulation of mitochondrial genome in different lineages.

Our novel discovery of a somatic mtDNA bottleneck specifically within the lymphoid lineage may play a role in the quality control of mitochondrial genomes, in parallel to the selection of immunoreceptor genes in the nuclear genome. Thus, a robust population of mtDNA may be crucial for lymphocyte-mediated immune responses. These findings provide new insight into immune degeneration and related diseases. The causing and the consequence of the somatic mtDNA genetic bottleneck require extensive efforts to explore.

## CODE AVAILABILITY

Code used for single cell data analysis and computational modeling are available at https://github.com/tangzhj/Bottleneck

## ACKNOWLEDGEMENTS

We thank Weiwei Zhai (Institute of Zoology, Chinese Academy of Sciences) for constructive comments and suggestions on the manuscript. This work was supported by National Natural Science Foundation of China (32070644) to J.X., Guangdong Basic and Applied Basic Research Foundation (2019A1515110387, 2019A1515110387) to J.X., the Natural Science Foundation of Guangdong (2021B1515020042) to Z.H., NIH grants RM1-HG007735 to H.Y.C. and China Postdoctoral Science Foundation (E12503) to Z.L. H.Y.C. is an Investigator of the Howard Hughes Medical Institute.

## AUTHOR CONTRIBUTIONS

J. X., Z.H and H.Y.C. designed and conceived the study. Z.T., B.C. and W.Z. collected and analyzed the data. Z.H. and Z.L. designed the dilution model and performed the simulations. J. X., Z.H., Z. T. and Z.L. wrote the manuscript with inputs from all authors. All authors read and approved the final manuscript.

## ONLINE METHODS

### Data collection

The mtscATAC-seq dataset generated through evaluation of hematopoietic and PBMCs was retrieved from a recent study evaluating samples from a healthy 47-year-old donor^17^. The mtscATAC-seq dataset from human bone marrow from 25-year-old healthy donor was obtained from Mimitou et.al^18^. The scATAC-seq data from CD4^+^ T cells were obtained from the study published by Satpathy et al.^37^. The scATAC-seq dataset for hematopoietic stem cells (HSCs), multi-potent progenitors (MPPs), lymphoid-primed multipotent progenitors (LMPPs), common lymphoid progenitors (CLPs), common myeloid progenitors (CMPs), granulocyte-macrophage progenitors (GMPs) and plasmacytoid dendritic cells (pDCs) derived from CD34^+^bone marrow was obtained from Buenrostro et al.^38^ (**Extended Data Fig. 2**). The scRNA-seq dataset generated from an evaluation of healthy CD34^+^ PBMCs, bone marrow mononuclear cells (BMMCs) and total PBMCs was downloaded from the study published by Granja et al.^39^ These datasets were used to analyze mtDNA replication and gene transcription (**Methods**). The scRNA-seq dataset of 70 effector memory T cells (Tem cells), 70 central memory T cells (Tcm cells) and 142 CD4^+^ regulatory T cells (Treg cells) from healthy human colon tissue were downloaded from Array Express (E-MTAB-6072)^40^. Detailed information on data resources is provided in **Supplementary Table 1**.

### Single-cell (sc)ATAC-seq data pre-processing and annotation of the cell populations

Raw data from GSE142745 were processed with Cell Ranger ATAC (version 2.0.3; 10x Genomics, https://www.10xgenomics.com/products/single-cell-atac) with default parameters. Reads were aligned to the reference hg19 human genome (https://support.10xgenomics.com/single-cell-atac/software/downloads/latest). In each cell, 40% of fragments overlapping a compendium of DNase hypersensitivity peaks and 1,000 unique nuclear fragments were filtered. From the output of the Cell Ranger Software calls, we performed a computational annotation of the cell types on the basis of chromatin accessibility. Clustering and gene activity scores were determined through standard processing via ArchR^41^. Clustering was performed with the “addClusters” and “addUMAP” functions (resolution=0.8, neighbors=10, minDist=0.1). To identify marker genes according to gene scores, we used the “getMarkerFeatures” function with useMatrix “GeneScoreMatrix” and generated a reproducible peak set in ArchR by using the “addReproduciblePeakSet” function. By default, ArchR attempts to identify peaks by using the MACS2 algorithm^42^. Because common cell markers are sometimes not suitable for classification with “GeneScoreMatrix”, we used enhancer accessibility to define the cell type. For example, we identified myeloid cells according to the unique accessibility of enhancers at +85 kb and +87 kb in the interferon regulatory factor (*IRF8*) locus. Plasmacytoid dendritic cells (pDCs) were identified on the basis of the unique accessibility of +54 kb and +56 kb enhancers, as described by Satpathy et al.^37^. Furthermore, to label scATAC-seq clusters with scRNA-seq information, we used the “addGeneIntegrationMatrix” function, which integrates scATAC-seq with scRNA-seq. Specific marker genes used to identify individual cell types in scATAC-seq datasets of healthy CD34^+^ hematopoietic cells and PBMCs are documented in **Supplementary Table 2**.

### Mitochondrial DNA variants identified in single-cell ATAC-seq datasets

Paired-end raw reads from each sample were aligned to the human reference genome (hg19) with Cell Ranger ATAC after adapter sequences were trimmed. First, the reads mapped to multiple sites or the nuclear genome, and duplicates were also removed. The remaining reads were realigned to correct the potential mapping errors around indels according to the process from GATK^43^. Bam files for each cell type were merged to identified germline mtDNA variants (bulk VAF >90%). Variants with VAF >90% shared among more than 90% cells were also considered germline mutations. Then mtDNA variants were called for each individual cell with VarScan2^44^ with “--min-var-freq 0.01” and “--min-reads2 2”. To identify high confidence somatic variants in single cell, the following filter steps were applied.

First, the germline mutations identified in the merged bam file were removed. Second, the following sites were explicitly removed because of the large numbers of homopolymers in the revised Cambridge Reference Sequence (rCRS) and sequencing errors in the reference genome^13^:

Misalignment due to ACCCCCCCTCCCCC (rCRS 302–315), including

302A/C, 309C/T, 311C/T, 312C/T, 313C/T and 316G/C;

Misalignment due to GCACACACACACC (rCRS 513–525), including

514C/A, 515A/G, 523A/C and 524C/G;

Misalignment due to 3107N in ACNTT (rCRS 3105–3109), including

3106C/A, 3109T/C and 3110C/A.

Third, sequencing errors can significantly affect the identification of somatic variants. Therefore, sequencing errors known to be associated with a high error rate according to Illumina NextSeq and sequence errors (G→T and C→A) from DNA damage were removed.

Fourth, strand balance was required for confident somatic variants. For the given variant site, we required the reads mapped to the forward strand to be above 30% but below 70% of the total mapped reads for the variant allele.

Variants that passed the multiple filter steps were merged from all individual cells as the final somatic variants. If the variant was sufficiently confident in any given cell, the variant allele frequency was re-counted in all individual cells within the same cell type, without any other constraints.

### Single-cell RNA-seq data processing and cell-type annotation

Downstream analysis of scRNA-seq dataset was performed with Seurat^45^ (version 3.2.2; https://satijalab.org/seurat). The following bioinformatic analyses were performed in R software (version 3.6.0; https://www.r-project.org) with default settings unless otherwise stated. Cells with <200 or >2,500 detected genes or with >5% mitochondrial DNA were eliminated from further consideration. Normalization was applied with the MAGIC package (version 2.0.3)^46^ by following the Seurat v3 workflow. We next calculated a subset of features that exhibited high cell-to-cell variability by using the “FindVariableFeatures” function and identified 2,000 specific features. Clusters were identified with the “Find-Neighbors” and “FindClusters” functions in Seurat with 45 principal components (PCs) and a resolution of 0.3. The results were annotated to include differential expression of cell type-specific marker genes. Uniform Manifold Approximation and Projection for Dimension Reduction (UMAP) dimensionality reduction was performed with the “RunUMAP” function in Seurat, with 45 PCs and other default parameters. The expression of cell type-specific marker genes in PBMCs and BMMCs is shown in **Supplementary Table 3**. We referred to the information and classifications recorded in GSE139369 from the GEO Database to guide our cell type annotations **(Supplementary Table 3)**.

### Pseudo-time analysis

To construct single-cell differentiation trajectories with scRNA-seq data from HSCs to B cells, we performed a pseudo-time analysis with the Monocle method^47-49^. First, we subdivided scRNA-seq data according to the annotated cell populations revealed by Seurat clustering analysis, according to the common pipeline (http://cole-trapnell-lab.github.io/monocle-release/monocle3/). Re-clustering of selected cell populations was again performed with the “RunUMAP” function. Pseudo-time analysis was conducted on these newly generated clusters with Monocle v3. We delineated expression patterns of G1/S phase-specific and mtDNA replication-related genes along a pseudo-timeline. G1/S phase-specific genes were identified according to a previously annotated list^50^ (**Extended Data Fig. 3d**).

### Mitochondrial DNA variants identified from single-cell RNA-seq data

Mitochondrial DNA variants from single-cell RNA-seq data were processed in the same manner as mtDNA variants from scATAC-seq, with several modifications. Briefly, we used STAR^51^ to align reads to the human reference genome (hg19) and to obtain bam files. Germline mutations and mtDNA variants in individual cells were filtered and called in the same manner.

### Allele frequency spectrum

The allele frequency (heteroplasmic ratio) of each mutation were calculated in each cell and the number of mutations fall in each frequency bin (from 0∼1) were counted for each cell types. Somatic mutations arose in the early development stage, which had been fixed in the progenitor cells, were further excluded for the ASF analysis in the mtscATAC-seq from BMMCs.

### Annotation of mitochondria DNA mutations and calculation of non-synonymous/synonymous mutation rates (dN/dS)

The mitochondrial variants were annotated with ANNOVAR^52^. The annotated variants comprised mutations in loops, tRNA, rRNA and mRNA coding regions, including non-synonymous (NS) and synonymous (SY) substitutions according to the variant location (**Extended Data Fig. 5a** and **Extended Data Fig. 6a**). Coding sequences (CDS) within the mitochondrial genome were evaluated with Phylogenetic Analysis of Maximum Likelihood (PAML) to identify all possible synonymous (defined as S) and nonsynonymous (defined as N) substitutions in the human mitochondrial genome^53^. On the basis of ANNOVAR’s annotations, we identified all observed synonymous (defined as s) and nonsynonymous substitutions (defined as n). The non-synonymous mutation rate (dN)=n/N and the synonymous mutation rate (dS)=s/S, responses to positive, neutral, or negative selection pressure, can be determined by the dN/dS ratio.

### Computational modeling of the mitochondrial genetic bottleneck

We used the Wright-Fisher model from population genetics to depict the accumulation of mutations and the dynamic frequency of heteroplasmic alleles in mtDNA during lymphoid cell divisions. The Wright-Fisher model assumes discrete generations and random sampling of individuals from the current generation without replacement by reproduction in the following generation. This model has been widely used to model the mtDNA population dynamics in both germline cells and somatic cells, including those that are neoplastic^23,54^. Because normal somatic cells typically contain 100– 1,000 copies of mtDNA, we used n=500 as the baseline copy number in our model^22^. Results from the scATAC dataset revealed that the relative copy number of mtDNA in NK cells was approximately 60% that detected in B or T cells (**Fig. 3a**); thus, 300 (500×0.6) was used as the baseline mtDNA copy number for the NK lymphocyte cohort. We modeled the lymphoid development from lymphoid-primed multipotent progenitor (LMPP) cells, which are the common progenitor cells for all lymphocytes, B, T and NK cells. To model the dilution-based genetic bottleneck, we introduced a dilution rate *α*, which denotes the fraction of mtDNA molecules in each cell that undergo replication within a single cell cycle, and *T*_*d*_, which denotes the time of the diluting process. After *T*_*d*_ cell divisions from LMPP, the mtDNA copy number in each cell type rapidly recovers to the baseline level. The minimal mtDNA copy number through the bottleneck can be computed by:

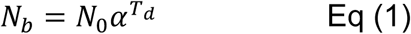

where *N*_*0*_ is the initial number of mtDNA copies. The total number of cell divisions required for the transition from an LMPP to a mature lymphocyte is denoted *T*_*a*_. The mutation rate at each site within the mitochondrial genome per cell division is denoted *μ*, which has been estimated to be 10^−8^–10^−7^ mutations per site for somatic cells^23,55^. Thus, the mutation rate for the entire mitochondrial genome during each cell division event will be *u* = *μ* × *L*, where *L*=16,569 base pairs (bp), representing the number of potential sites within the mitochondrial DNA length.

During each cell division, the number of somatic mutations acquired per mitochondrial genome follows a Poisson distribution with a mean of *u*. Thus, the probability that k mutations occurred in each cell division is as follows:

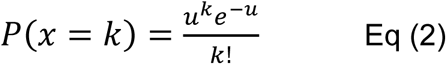

### Computational inference of parameters by approximate Bayesian computation

We used the framework of approximate Bayesian computation (ABC) for parameter inference in our computational model of somatic mtDNA population dynamics on the basis of the dilution rate *α*, the dilution time course *T*_*d*_ and the total number of cell divisions *T*_*a*_. The minimal mtDNA copy number in each cell can be computed as described by Eq (1) when values for *α* and *T*_*d are*_ available. The prior uniform distributions used for sampling *α, T*_*d*_ and *T*_*a*_, were *α∼U*(0,1) *T*_*d*_*∼U(*0, 30) and *T*_*a*_*∼U*(10, 40). avoid extinction (i.e., minimal mtDNA copy number=0), only the sampled parameter values ensuring 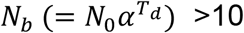 were retained. We used a version of ABC based on the acceptance-rejection algorithm^56^to estimate posterior probability distributions for the parameters of interest (i.e., ***θ*** *[*α*, T*_*d*_, *T*_*a*_]. We used 19 summary statistics (***S***), which included the mtDNA mutation count in each VAF bin as step=0.05 from VAF=0.05 to 1 to fit the simulated to the observed data. The ABC version of rejection sampling is as follows:

For *i*=1 to *K* simulations:

1. Sample parameters **θ′** from the prior distribution *π*(**θ**)
2. Simulate data (**D′**) with the sampled parameters (**θ′**) and summarize **D′** as summary statistics (**S′**).
3. Accept **θ′** if *d*(**S′, S**)**<***ε*, for a given tolerance rate *ε*, where *d*(**S′, S**) is a measure of the Euclidean distance between **S′** and **S**
4. Return to step 1.

With this scheme, we approximated the posterior distribution by *P*(***θ***|*d*(**S**′, **S**)<*ε*). We used a common variation in ABC^57,58^ in which, rather than using a fixed threshold, *ε*, we sorted all calculated *K* distances by *d*(**S′, S**) (see step 3 above) and accepted the ***θ′*** that generated the smallest 100×*η* percentage distances. We used *K*=10^6^ and *η*=0.001 so that the posterior distribution was composed of 10^6^×0.001=1,000 data points. We ran the ABC inference procedures for two mutation rates (*μ* =10^−8^ and 10^−7^) and performed model selection (**Extended Data Fig. 4**). The mutation rate *μ* =10^−7^ fitted the data better in all cell types and thus was used for the computational inference. The ABC procedure was performed with the R package *abc*^59^.

## Figures and Legends

**Extended Data Fig. 1.**
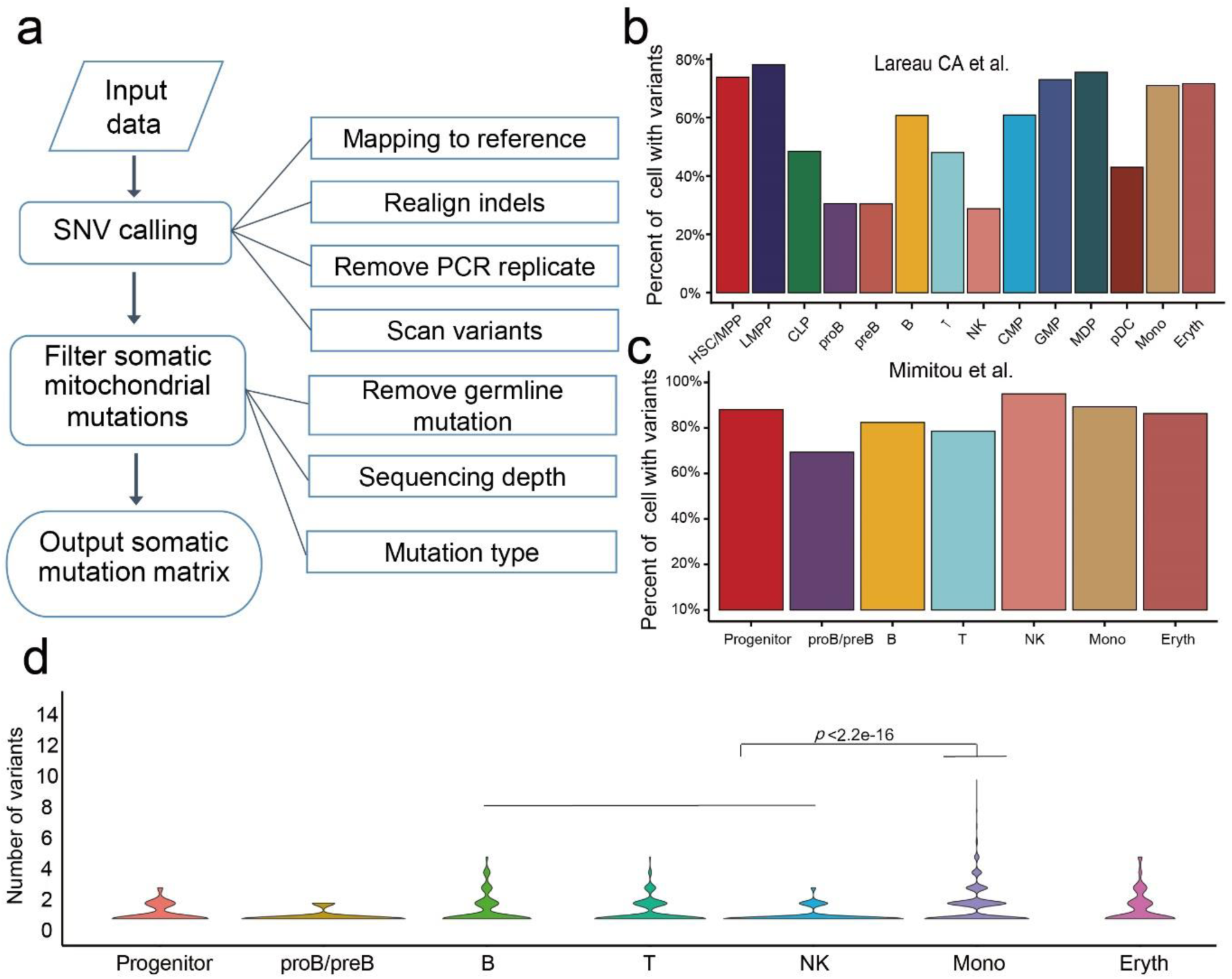
Detection of somatic mitochondrial mutations in single PBMCs with scATAC-seq data or scRNA-seq data. **(a)** Schematic of mtDNA mutation calling with scATAC-seq (including mtscATAC-seq) or scRNA-seq data. **(b)** Percentage of cells with at least one somatic mtDNA mutation detected in individual cells for each cell type in the mtscATAC-seq data from Lareau et al. **(c)** Percentage of cells with at least one somatic mtDNA mutation detected in individual cells for each cell type in the mtscATAC-seq data from Mimitou et al. **(d)** Violin plot showing the number of somatic mtDNA variants per cell for various cell types; *p* values based on a two-sided Wilcoxon rank-sum test are as shown.

**Extended Data Fig. 2.**
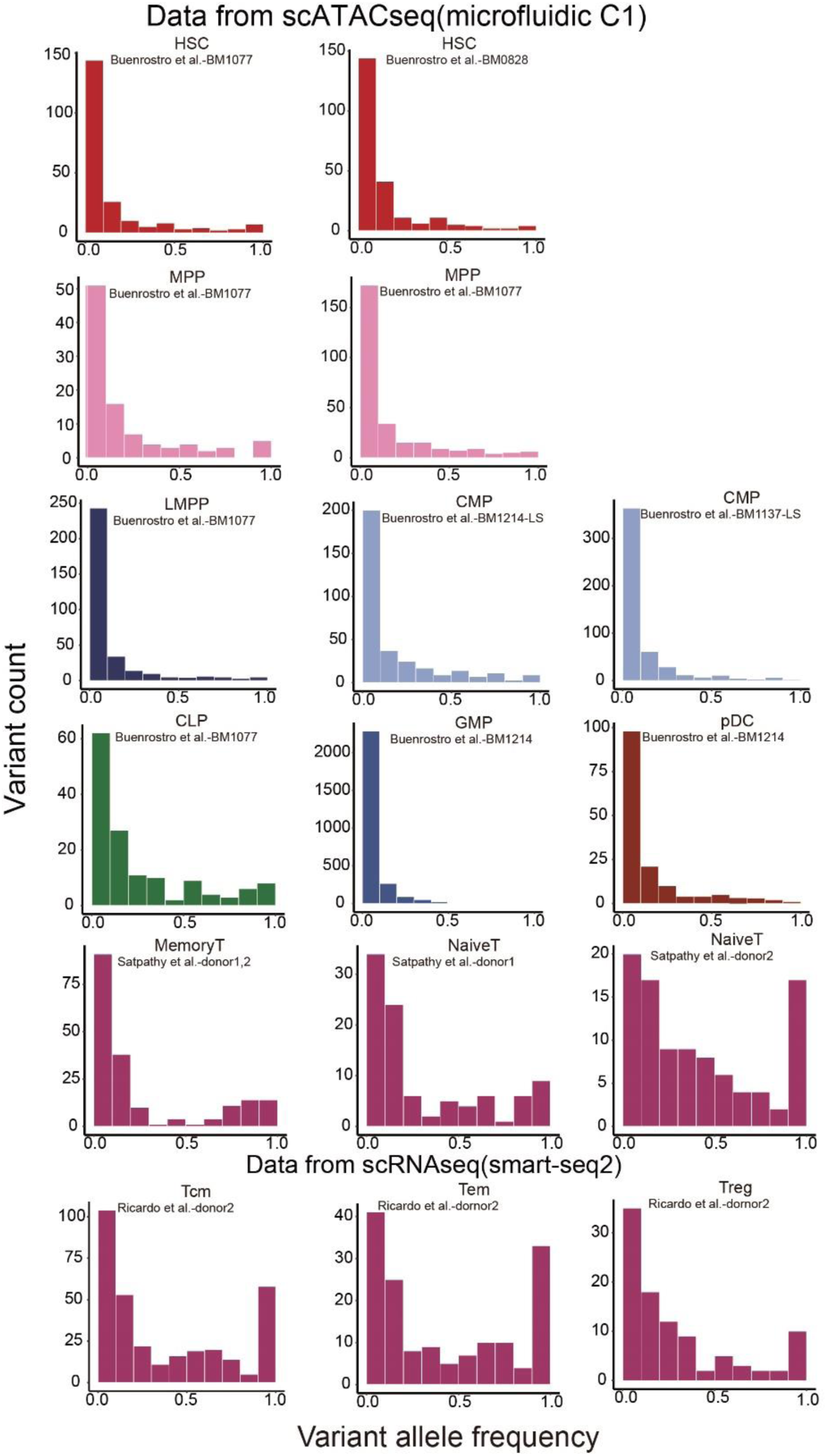
Allele frequency spectrum of somatic mtDNA mutations for different hematopoietic cell types, on the basis of independent scATAC datasets (Buenrostro et al. and Satpathy et al.) and an scRNA-seq dataset (Ricardo et al.).

**Extended Data Fig. 3.**
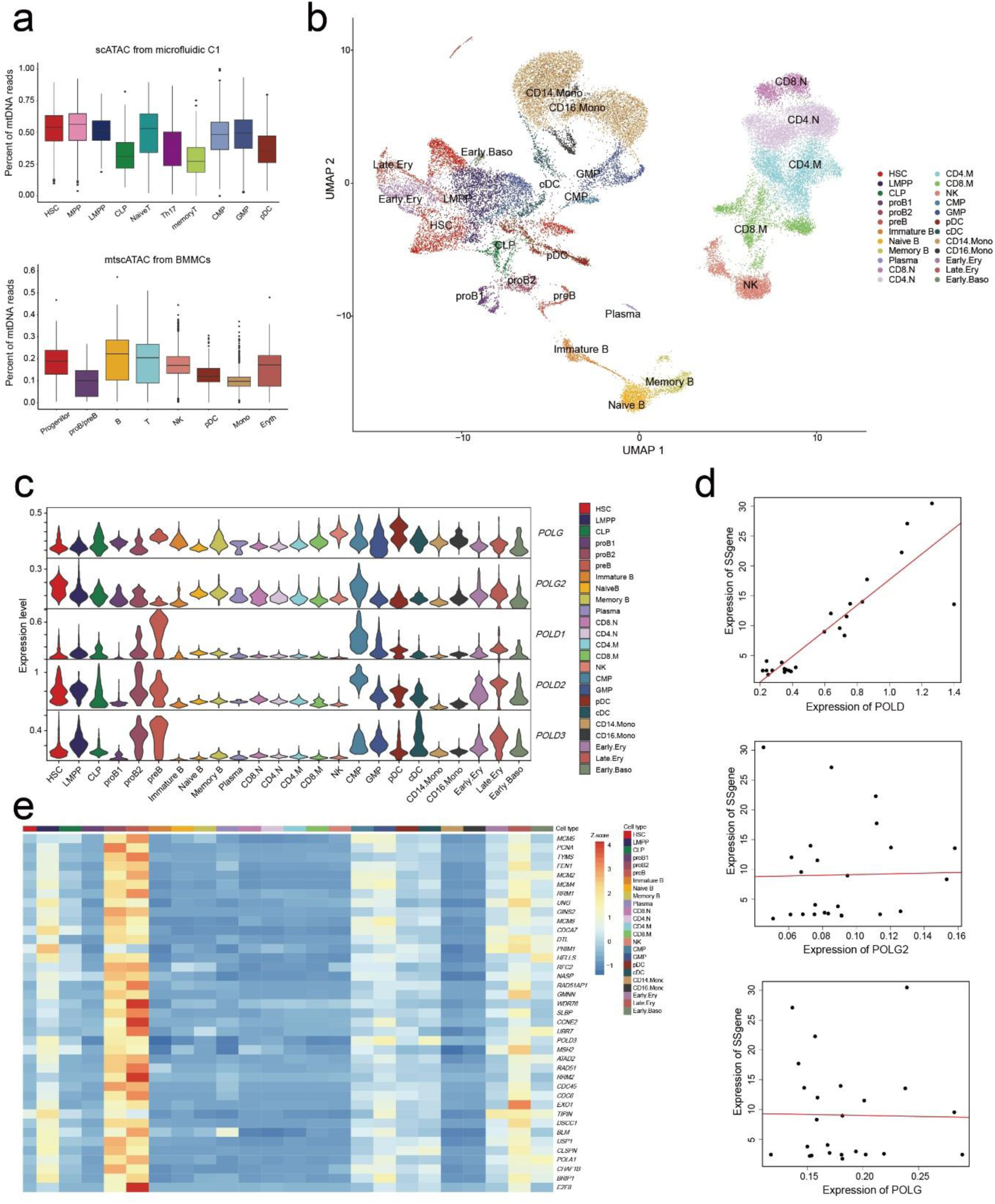
Gene expression of G1/S phase-specific genes in scRNA-seq data. **(a)** Relative mtDNA copies were measured by the percentage of the sequencing reads mapped to the mitochondrial genome out of the total number of reads for each cell types. **(b)** UMAP projection of PBMCs, BBMCs and CD34^+^ PBMCs with scRNA-seq data. Dots represent individual cells colored by cell types. **(c)** Violin plots showing the expression of mitochondrial DNA polymerase γ (*POLG*) and its binding subunit (*POLG2*) and nuclear DNA replication polymerase genes (*POLD1–3*) from scRNA-seq data. **(d)** Scatter plot showing the correlation of the gene expression of *POLD* (*POLD1–3*), *POLG* and *POLG2* with G1/S phase-specific genes (SSgene). **(e)** Heat map showing the expression of 39 G1/S phase-specific genes in 24 cell types.

**Extended Data Fig. 4.**
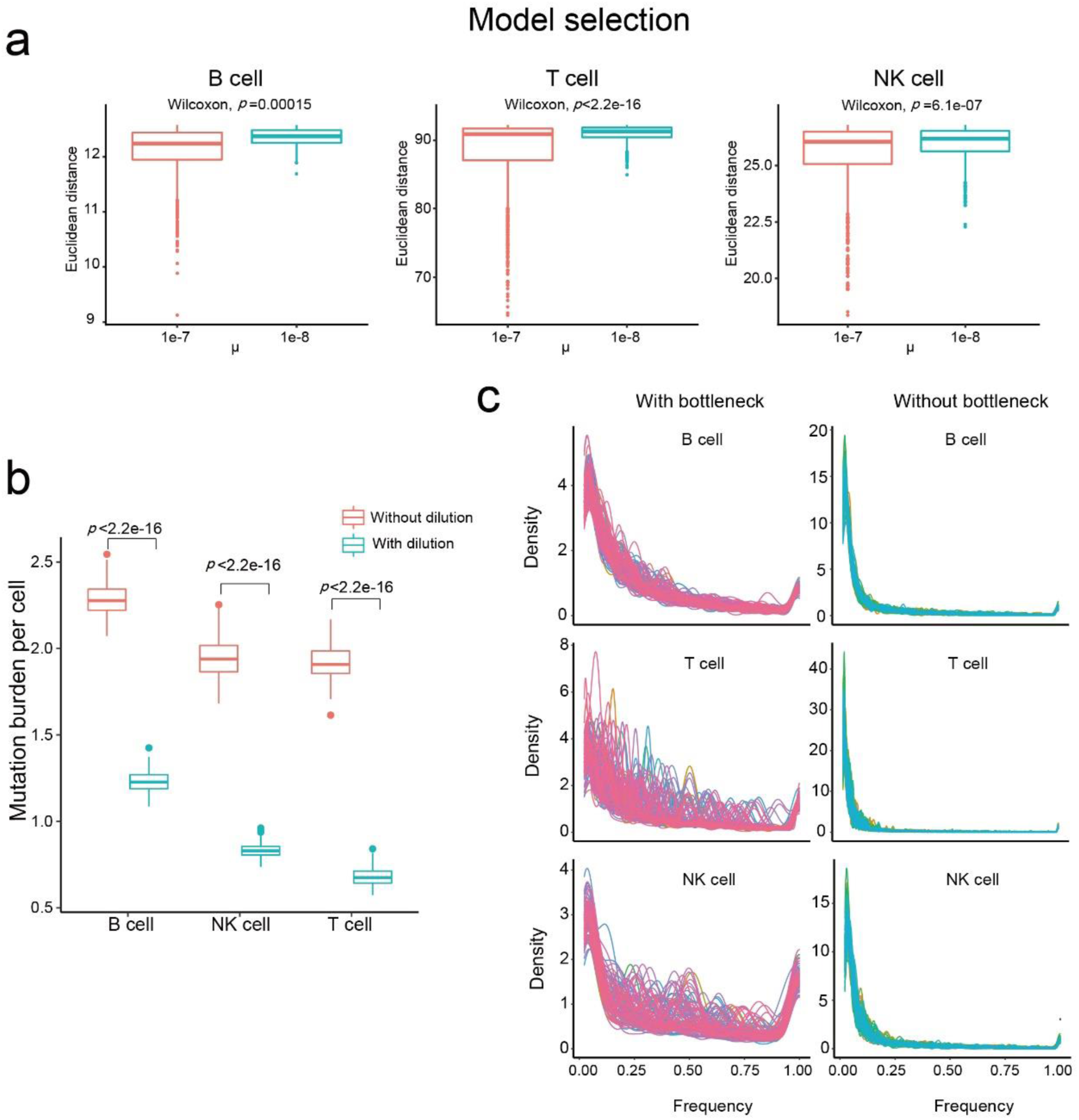
Details regarding the parameter inference for the dilution model of the mitochondrial genetic bottleneck. **(a)** Model selection with respect to the per-site mutation rate *μ*. We ran the ABC inference procedures for two mutation rates *μ* =10^−8^ and 10^−7^, and *μ* =10^−7^ fitted the data better (smaller Euclidean distance between simulated and observed summary statistics) in all cell types and thus was used for the parameter inference. (**b**) Simulations under the dilution model of the mitochondrial genetic bottleneck with the ABC-estimated parameter values recapitulated the lower mutation burden in B, T and NK cells, as compared with simulations without a mitochondrial genetic bottleneck. (**c**) Simulations (100 times) with inferred parameters from the dilution model under conditions with or without a mitochondrial genetic bottleneck. Each curve represents one simulation.

**Extended Data Fig. 5.**
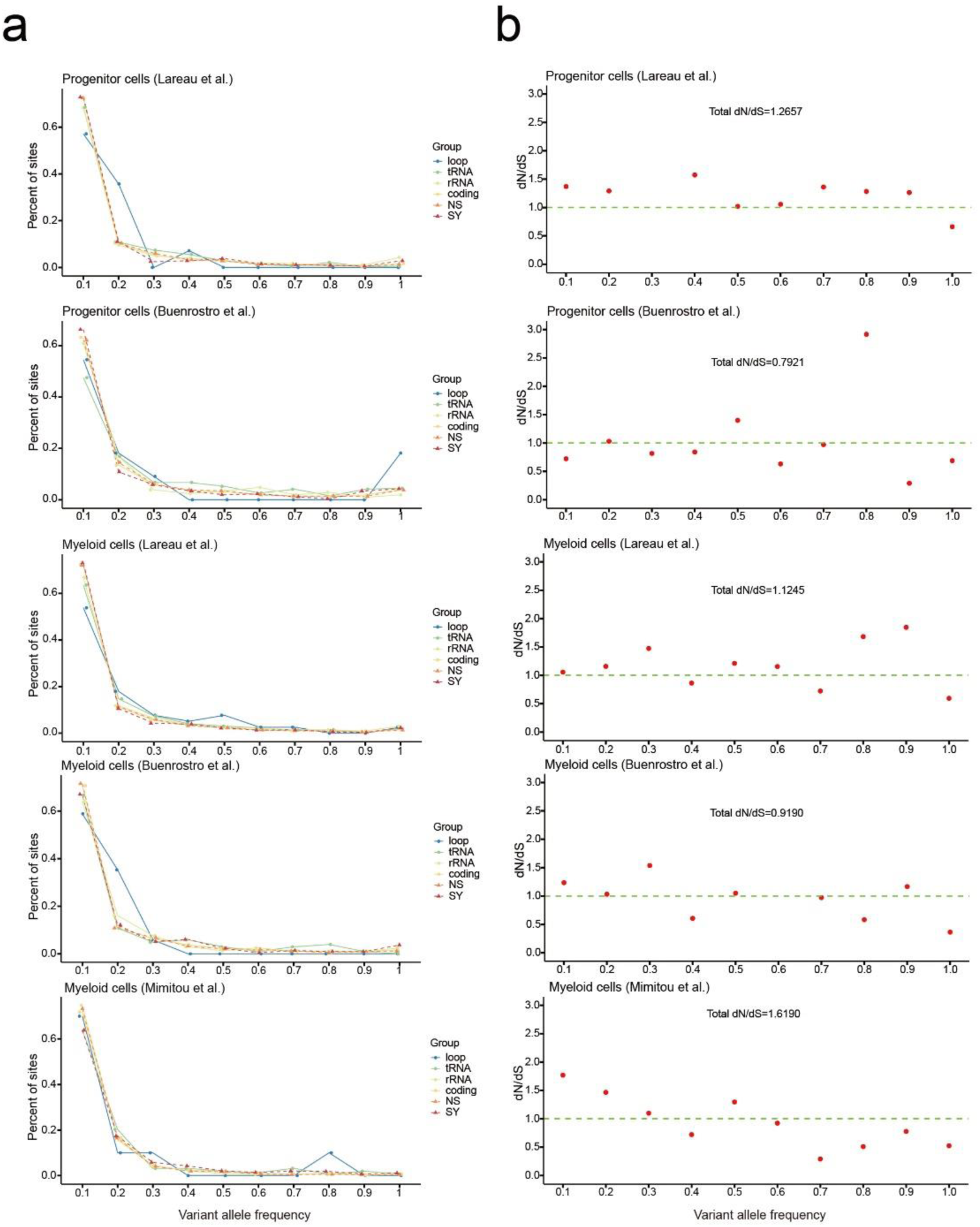
Allele frequency spectrum of somatic mtDNA mutations for different types and dN/dS in the progenitor and myeloid lineages. (**a**) Distribution of the VAF for mutations in different mtDNA genomic regions or types in progenitor and myeloid cells. The color code corresponds to mtDNA genomic regions or mutation types, annotated as loop, tRNA, rRNA, coding (coding region), NS (non-synonymous) and SY (synonymous). (**b**) The dN**/**dS ratio (y-axis) for mutations in different VAF bins (x-axis).

**Extended Data Fig. 6.**
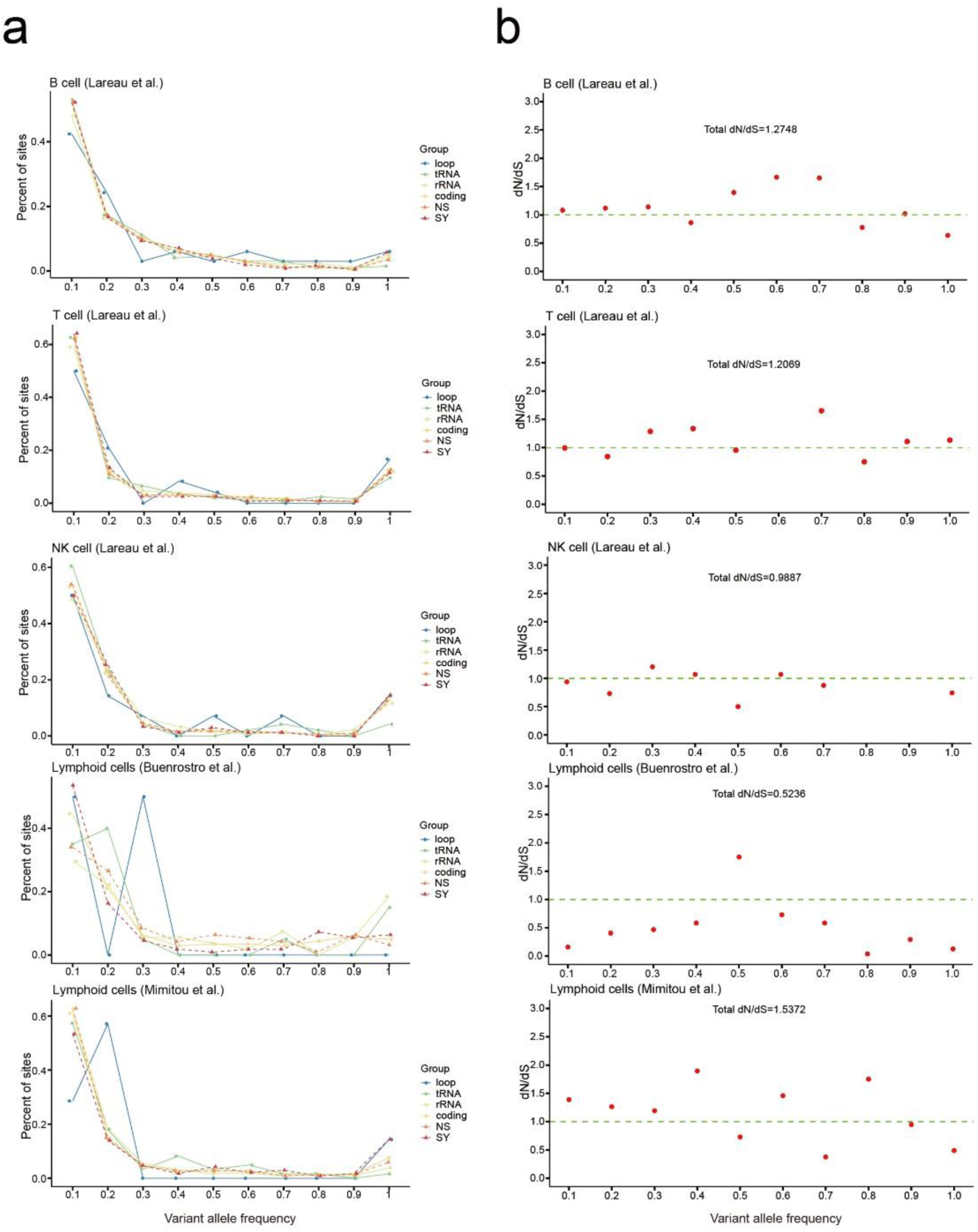
Allele frequency spectrum of somatic mtDNA mutations for different types and dN/dS in the lymphoid lineage. (**a**) Distribution of VAF for mutations in different mtDNA genomic regions in lymphoid cells (B, T and NK). The color code corresponds to mtDNA genomic regions or mutation types, annotated as loop, tRNA, rRNA, coding (coding region), NS (non-synonymous) and SY (synonymous). (**b**) dN**/**dS ratio (y-axis) for mutations in different VAF bins (x-axis).

**Supplementary Table 1.**
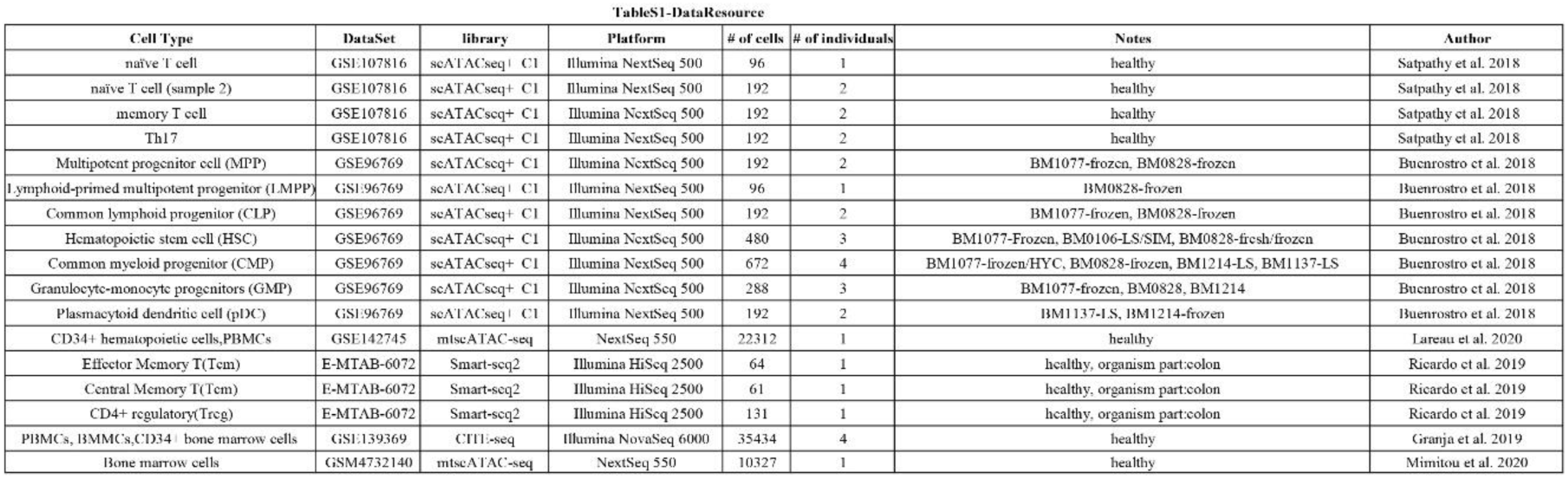

**Supplementary Table 2.**
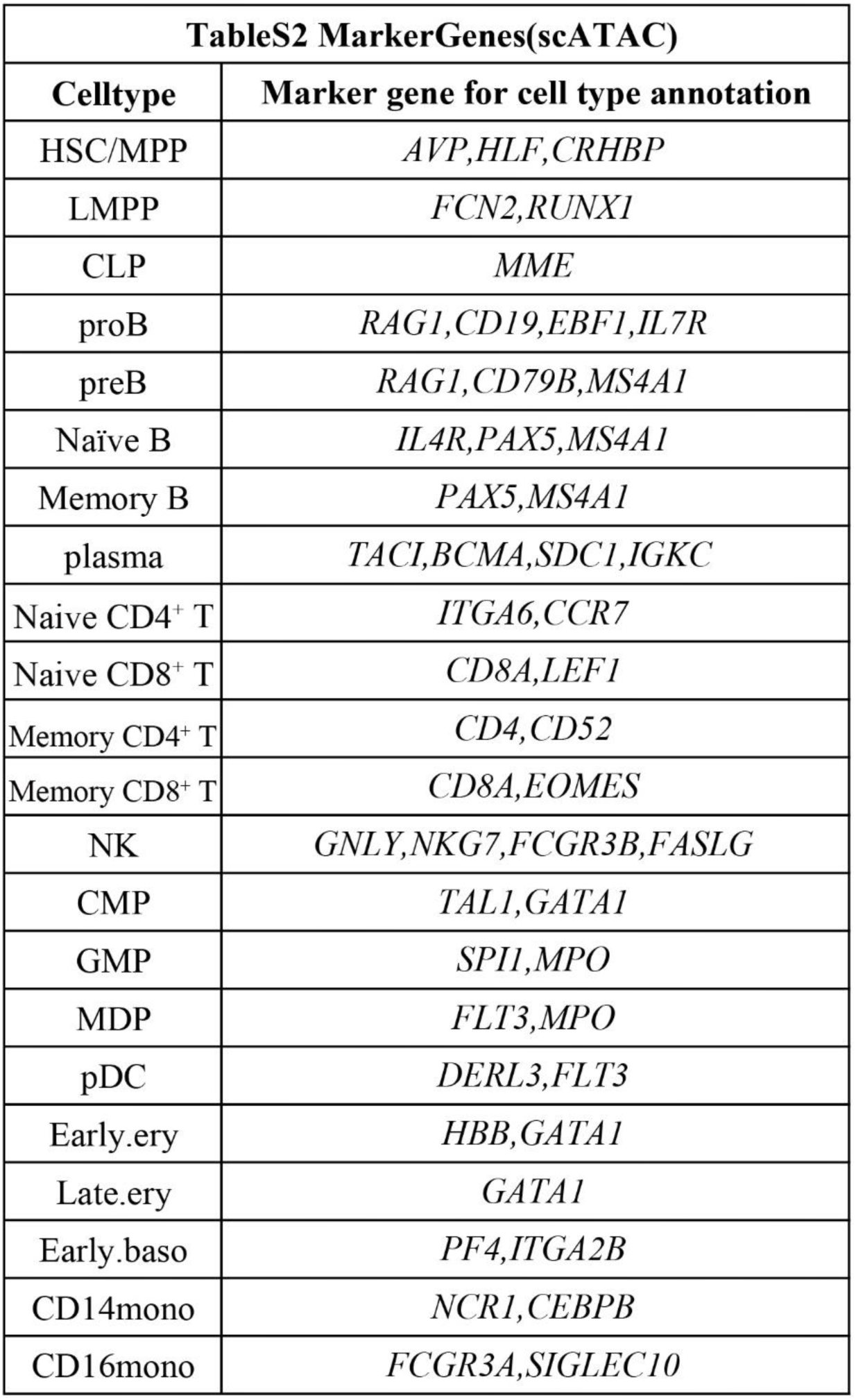

**Supplementary Table 3.**
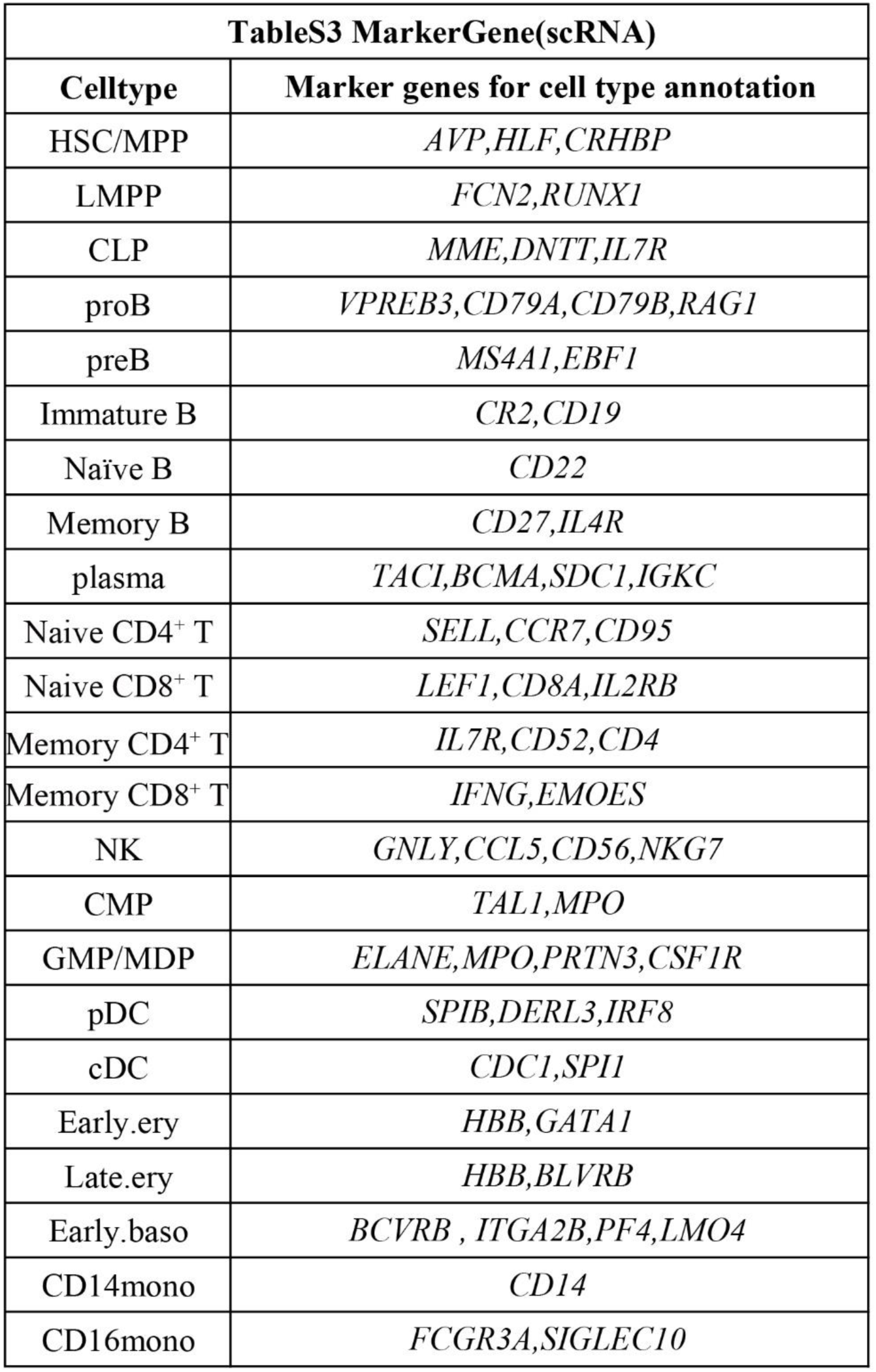

